# Transcriptomic mapping of the metzincin landscape in human trophoblasts

**DOI:** 10.1101/2022.02.15.480614

**Authors:** Jasmin Wächter, Matthew J Shannon, Barbara Castellana, Jennet Baltayeva, Alexander G. Beristain

**Affiliations:** The British Columbia Children’s Hospital Research Institute, Vancouver, Canada; Department of Obstetrics & Gynecology, The University of British Columbia, Vancouver, Canada

**Keywords:** Placenta, trophoblast, protease, metzincin protease, matrix metalloprotease, progenitor, single cell RNA sequencing, differentiation, organoids, ADAM metalloprotease With Thrombospondin Type 1 Motif 6, Fibrillin 2

## Abstract

The metzincin family of metalloproteases coordinates cell and tissue developmental processes through regulation of growth factor availability, receptor signaling, and cell-cell/cell-matrix adhesion. During placental development, while distinct roles for metzincin proteases in controlling specific trophoblast functions have been described, a comprehensive assessment of metzincins during discrete stages of trophoblast differentiation has yet to be performed. Here we provide a comprehensive single cell transcriptomic resource of metzincin protease expression in diverse states of human trophoblasts from first trimester placental and decidual tissues. In the 8 distinct trophoblasts states categorized [four progenitor cytotrophoblast (CTB), one syncytiotrophoblast precursor (SCTp), two column CTB (cCTB), and one extravillous trophoblast (EVT) state], we identified 24 metzincin genes. These included 12 adamalysins, 2 pappalysins, 3 astacins and 7 matrixins. Cell trajectory modeling shows that expression of most (19/24) metzincins increases across CTB to EVT differentiation, though select proteases also increase as CTB fuse into syncytiotrophoblast. Within the CTB niche, single-cell velocity ordering identified 11 metzincins (*ADAM10, -17, MMP14, -15, -19, -23B, ADAMTS1, -6, -19, TLL-1, -2*) expressed in progenitors proximal to the predicted origin. Analysis of metzincin-substrate interactions within the CTB niche revealed ∼150 substrates and binding partners, including *FBN2* as an *ADAMTS6*-specific substrate preferentially expressed in trophoblast progenitors. Together, this work characterizes the metzincin transcriptomic landscape in human first trimester trophoblasts and establishes insight into the roles specific proteases perform within distinct trophoblast niches and across differentiation. This resource serves as a guide for future investigations into the roles of metzincin proteases in human placental development.

**Summary Statement:** Single cell RNA sequencing characterizes the expression of multiple metzincin proteases within first trimester placental trophoblasts. Examination of protease-substrate interactions within cytotrophoblasts identifies potential interactions between ADAMTS6 and FBN2.

**Highlights:**

- Single cell RNA sequencing identifies 24 distinct metzincin proteases expressed in human first trimester trophoblasts
- Lineage trajectory modelling shows that metzincin genes are dynamic and likely control processes in progenitor, mid-point, and end-point states of trophoblast differentiation.
- ADAMTS6, and its putative substrate FBN2, localize specifically to progenitor trophoblasts

## INTRODUCTION

The placenta is a transient organ that develops during pregnancy and serves as a physiological link between the developing fetus and mother. Multiple processes are coordinated by the placenta, where cross-species similarities in function amongst placental mammals are generally conserved [1]. For example, the placenta facilitates nutrient and waste exchange between fetal and maternal systems, produces key hormones and growth factors required for pregnancy maintenance, and modulates the maternal immune system to tolerate the semi-allogeneic fetus and placenta [2,3]. Therefore, it may come as no surprise that abnormal placentation is a confounding feature of many pregnancy disorders [4–6]. Central to many of the placenta’s functions are the trophectoderm-derived trophoblasts.

In humans, trophoblast progenitors, termed cytotrophoblast (CTB), differentiate along two major cell pathways: villous and extravillous [3,7]. CTB reside within the inner trophoblast layer of floating villi or proximal to areas of placental-uterine attachment. In floating villi, CTB proliferate and fuse with neighboring CTB to generate an overlying multinucleated structure called the syncytiotrophoblast (SCT), representing the main site of physiological exchange between fetal and maternal circulations and the major endocrine engine of the placenta [2]. Alternatively, CTB proximal to placental-uterine attachment points proliferate and expand to form multilayered columns of cells called column CTB (cCTB). As cCTB mature, epithelial characteristics (i.e., tight cell-cell adhesions) are gradually lost and replaced with mesenchymal-like features (i.e., reduced homotypic adherence, acquisition of mesenchymal molecular programs) [8,9]. At distal ends of anchoring columns, cCTB detach and invade into the maternal compartment where they are now considered terminally differentiated extravillous trophoblast (EVT) [3]. EVT are highly motile and invade/migrate into uterine arteries, veins, and lymphatics, and in doing so play central roles in coordinating uterine vascular remodeling and tempering the maternal immune system [10]. While these invasive features of EVT are not dissimilar from invasive/metastatic cancer cells, EVT invasion is a highly regulated process that is coordinated in part by soluble factors produced by cells of the decidualized endometrium.

Metzincin metalloproteases are a diverse family of proteases defined by the presence of a methionine residue at the active/regulatory site and the incorporation of a zinc ion within the active site that is necessary for enzymatic reactivity [11,12]. Metzincin metalloproteases are comprised of matrix metalloprotease (MMP), a disintegrin and metalloprotease (ADAM), ADAM with thrombospondin repeat (ADAMTS), astacins, snapalysin, serralysin, leishmanolysin and pappalysin subfamilies. The importance of metzincin proteases is underlined by key roles of MMPs in tissue morphogenesis (i.e. tadpole tale resorption; [13]) and their involvement in chronic diseases like arthritis and cancer [14,15]. Twenty-four MMPs, twenty-one ADAMs, nineteen ADAMTS, six astacins, and two pappalysins are present in the human genome, and together play diverse roles in extracellular matrix remodeling/degradation [16], receptor/ligand activation and availability [11], and the control of cell-cell and cell-matrix interactions [17].

The diverse biological processes regulated by metzincin metalloproteases suggest that they may play central roles in placentation, and in particular, trophoblast development and function. Indeed MMP-directed matrix degradation controls EVT invasion [18,19], while ADAM-controlled cell-matrix and cell-cell processes play roles in promoting anchoring villi outgrowth and EVT motility [20,21], as well as SCT formation [22]. More recently in other systems, the importance of metalloproteases in controlling the availability of soluble factors in defined cell niches suggests that protease-controlled events may regulate stem cell and lineage fate decisions during tissue/organ development [23,24]. However, it is not currently known if metzincin proteases play roles in CTB progenitor maintenance or downstream sub-lineage commitments.

Advances in single cell transcriptomics allow for unprecedented examination of cell heterogeneity, cellular hierarchy, and gene pathways central to stem cell maintenance and the control of differentiation. In this work, we use published single cell RNA sequencing (scRNA-seq) datasets [25,26] to define the metzincin protease landscape of human trophoblasts in early pregnancy. We show that human trophoblasts express 24 metzincin proteases. Bioinformatic cell lineage modeling describes metzincin protease kinetics during SCT and EVT differentiation, and importantly sheds light on the importance of six proteases (*ADAMTS6, ADAMTS19, ADAM10, ADAM17, MMP14*, and *MMP15*) as well as their putative substrates in the CTB progenitor niche. Together, this work defines the metalloprotease transcriptomic landscape in early human trophoblast development. Importantly, this work also provides detailed insight and a comprehensive resource for understanding the role of metzincin proteases in shaping the CTB progenitor niche.

## MATERIALS AND METHODS

### Patient recruitment and tissue collection

Placental tissues were obtained with approval from the Research Ethics Board on the use of human subjects, University of British Columbia (H13-00640). All samples were collected from pregnant individuals (19 to 39 years of age) providing written informed consent undergoing elective terminations of pregnancy at British Columbia’s Women’s Hospital, Vancouver, Canada. First trimester placental tissues (*n*=3) were collected from participating pregnant individuals (gestational ages ranging from 6–12 weeks) having confirmed viable pregnancies by ultrasound-measured fetal heartbeat. Patient clinical characteristics i.e., height and weight were additionally obtained to calculate body mass index (BMI: kg/m^2^). All consenting pregnant individuals provided self-reported information via questionnaire to having first-hand exposure to cigarette smoke or taken anti-inflammatory or anti-hypertensive medications during pregnancy. Patient and sample phenotype metadata are described in Supplemental Table 4.

### Placental tissue single cell RNA-seq library construction

#### Data Repository Integration

Droplet-based first trimester scRNA-seq decidual (n=4) and placental tissue (*n*=11) data was obtained from the public repository ArrayExpress (E-MTAB-6701) [25] and from the GEO repository (GSE174481) [26]. Data was integrated and sample specific metrics are summarized as in Shannon *et al*. [26].

### Single cell RNA-seq data analysis

Code used for all subsequent data processing and analyses are available at: https://github.com/JasminWaechter and https://github.com/MatthewJShannon.

#### Data pre-processing and quality control

A total of 50,790 single cells from maternal-fetal interface samples were sequenced and pre-processed using the Seurat R package (version 4.0.1) [27,28]. Quality control, cell cycle scoring, scaling, normalization, removal of doublets and integration was done as previously described in Shannon *et al*. [26].

#### Cell clustering, identification, and trophoblast sub-setting

Cell clustering, trophoblast identification, and subsequent filtering for trophoblasts was done identically as described in Shannon *et al*. [26]. Trophoblasts were subset and re-clustered in Seurat at a resolution of 0.375 using 50 principal components. Trophoblast projections were visualized as Uniform Manifold Approximation and Projections (UMAPs).

#### Protease Identification

Expression of all known metzincin proteases was examined in trophoblasts. Proteases were considered significantly expressed when either showing expression in 1% of all cells or 5% of cells in a specific cluster. Similar thresholds have been used elsewhere [29][30].

#### Pseudotime

The Monocle2 R package (version 2.18.0) [31–33] was used to explore the differentiation of progenitor CTB into specialized SCT and EVT. The trophoblast raw counts were loaded into Monocle2 for semi-supervised single cell ordering across a pseudotime vector, using *BCAM* and *EGFR* expression to denote cells at the origin and *HLA-G* and *ITGA5*, as well as *ERVFRD-1* and *ERVV-1* gene expression to indicate progress towards EVT and SCT, respectively. Branched expression analysis modelling (BEAM) was applied to observe the expression of the 24 identified proteases over pseudotime, progressing towards either EVT or SCT. Results were visualized using the “plot_cell_trajectory” and the “plot_genes_branched_pseudotime” functions. To further confirm trophoblast differentiation trajectories, the Monocle3 package version 1.1.0 [34] was used as described in Shannon *et al*. [26].

#### Lineage Trajectory

Identical to Shannon et al. [26], sample-specific count matrices of unspliced, spliced, and ambiguous RNA transcript abundances were generated using the Velocyto package [35] and merged with their corresponding 10X sample using velocyto.R (version 0.6). The final, processed trophoblast object and UMAP embeddings were exported to a Jupyter Notebook and the scVelo Python package [36] was used to read, prepare, and recover holistic trophoblast gene splicing kinetics using the 2000 most differentially expressed genes. Velocities were projected on top of the extracted UMAP embeddings generated in Seurat.

#### Differential gene expression

Differential expression (DEG) analyses were performed in Seurat using the “FindMarkers” function using a MAST [37] framework on the raw count data. Ident.1 was set to CTB1-4. Parameters included a minimum log fold-change difference between groups of “−INF”, a minimum gene detection of “−INF” in cells in each group, and a minimum difference in the fraction of detected genes within each group of “−INF”, allowing all genes to be compared. Results were visualized using the R package EnhancedVolcano (version 1.8.0), with a fold-change cut-off of >0.5 and a p value cut-off of <10e-4 as the thresholds for significance.

#### Protease – Substrate Analysis

The NicheNet package [38] was used to examine protease-substrate and protease-binding partner interactions. Additional interactions and substrates taken from the literature were added to the existing NicheNet-curated database. Their respective sources and experimental evidences are shown in Supplemental Table 3. To build the updated “ligand-target-matrix”, new data points were given a source weight of 1 to provide a strong contribution in the final model. Afterwards, data sources were aggregated using “construct_weighted_networks”. Source weights were optimized using “apply_hub_corrections” and finally, the “ligand-target matrix” was built using “construct_ligand_target_matrix”. For the analysis, both receiver and sender cells were denoted as CTB1-4. Proteases expressed in 20% of CTB were selected as “ligands”. Substrates and binding partners expressed in 20% of CTB were considered for the analysis. Several sources including gene regulatory information, cell signaling databases, phosphorylation network, kinase-substate information and indirect interaction were excluded from the protease-substrate interaction analysis. Information regarding exclusion of sources is described in Supplemental Table 2. The resulting protease-substrate interactions were visualised using the “vis_ligand_receptor_network” function. Literature curated substrates (blue) and interactions (lime green) are marked. Specific signaling pathways were extracted using “get_ligand_signaling_path”. All data sources were included for the aforementioned follow-up analysis. The percentage of cells in CTB 1-4 expressing each gene in the extracted signalling pathway was manually added as a colour scale (red).

#### Gene ontology

GO analysis of expressed genes encoding identified substrates and interactions was performed using the clusterProfiler R package (version 3.18.1) [39] function “enrichGO” and visualized using the “goplot” function. Specific FBN2 pathways were examined using the ReactomePA package (version 1.36.0) [40]. The “Elastic fibre formation” pathway was highlighted.

### Immunofluorescence microscopy

Placental villi (6-12 weeks’ gestation; *n*=3) were fixed in 2% paraformaldehyde overnight at 4°C. Tissues were paraffin embedded and sectioned at 6 µm onto glass slides. Immunofluorescence was performed as previously described [41]. Briefly, placental tissues underwent antigen retrieval by heating slides in a microwave for 5 × 2-minute intervals in 10mM citrate buffer (pH 6.0). Sections were incubated with sodium borohydride for 5 minutes at room temperature (RT), followed by Triton X-100 permeabilization for 5 minutes, RT. Slides were blocked in 5% normal goat serum/0.1% saponin for 1 hr, RT, and incubated with combinations of the indicated antibodies overnight at 4 °C: mouse monoclonal anti-HLA-G (clone 4H84; 1:100; Exbio); rabbit monoclonal anti-cytokeratin 7 (clone SP52; 1:50; Ventana Medical Systems), rabbit polyclonal anti-EGFR (D38B1) (1:50; clone: 4267; Cell signalling), rabbit polyclonal anti-ADAM-17 (1:100; ab19027; Abcam), mouse monoclonal anti-hCG beta (clone 5H4-E2; 1:100; Abcam), purified rabbit polyclonal isotype ctrl (1:100; clone: 910801; Biolegend)

Following overnight incubation, sections and coverslips were washed with PBS and incubated with Alexa Fluor goat anti-rabbit 568 and goat anti-mouse 488 conjugated secondary antibodies (Life Technologies, Carlsbad, CA) for 1 hr at RT, washed in PBS and mounted using ProLong Gold mounting media containing DAPI (Life Technologies). Slides were imaged with an AxioObserver inverted microscope (Car Zeiss, Jena, Germany) using 20X Plan-Apochromat/0.80NA or 40X Plan-Apochromat/1.4NA objectives (Carl Zeiss). An ApoTome .2 structured illumination device (Carl Zeiss) set at full Z-stack mode and 5 phase images were used for image acquisition. Patient and sample phenotype metadata are described in Supplemental Table 4.

### Mean fluorescent intensity measurements

Ten snapshots of HLA-G expressing anchoring villi were taken for each placenta (*n*=3). Analyses were performed in ImageJ (version 2.0.0). For each image, background was subtracted. Sections were visually divided into proximal column, medial column, and distal column based on HLA-G expression intensity. Mean values from each section were measured. These values were then divided by the mean intensity value of the whole image.

### RNAscope

Placental villi (6-12 weeks’ gestation; *n*=2) were fixed in 2% paraformaldehyde overnight at 4°C. Tissues were paraffin embedded and sectioned at 6 µm onto glass slides. The RNAscope 2.5HD Duplex Assay (ACD Bio) was performed according to manufacturer’s instructions. The Hs-FBN2-C2-*Homo sapiens* fibrillin 2 mRNA (Cat#1075921-C2, ACD bio) probe and the Hs-ADAMTS6-*Homo sapiens* ADAM metallopeptidase with thrombospondin type 1 motif 6 mRNA (Cat#814831) were used. Slides were imaged with an Olympus upright microscope (Olympus corporation, Japan) using 20X Plan-Apochromat/0.80NA or 40X Plan-Apochromat/1.4NA objectives (Olympus Corporation, Japan). Patient and sample phenotype metadata are described in Supplemental Table 4.

### Statistical analysis

Mean fluorescent intensity data are reported as median values with standard deviations. All calculations were carried out using R software. Measurements were examined for normality using the Shapiro-Wilks test. The homogeneity of variance was examined using the Levene’s Test. Outliers were removed. One-way ANOVA followed by Tukey post-test were performed for all other measurements. The differences were accepted as significant at *P* < 0.05.

## RESULTS

### The metzincin protease landscape within transcriptionally-defined trophoblast states

Single cell transcriptomics (scRNA-seq) enables the identification of cellular heterogeneity and hierarchy within complex tissues and organs. Moreover, scRNA-seq allows for the comprehensive analysis of gene profiles at single cell resolution. Recently, we established scRNA-seq libraries from seven unique first trimester chorionic villus tissues [26] and integrated these data with a publicly available scRNA-seq dataset consisting of chorionic villus (*n*=4) and decidual (*n*=4) specimens [25]. Using these published datasets, we set out to comprehensively examine the metzincin protease transcriptomic landscape in human trophoblasts. Implementing our previously described single cell dataset filtering and analysis pipeline [26], 7,798 trophoblasts originating from chorionic villi and decidua were subset from non-trophoblast cells and resolved into transcriptionally-defined clusters that identify unique trophoblast states (Fig. 1A). Recapitulating our previous findings [26], eight cell states were resolved, and by referencing known trophoblast marker genes, four CTB (CTB1-4; *TEAD*^+^, *EGFR*^+^), one syncytiotrophoblast precursor (SCTp; *ERVFRD-1*^+^), two column CTB (cCTB1-2; *ITGA5*^+^, *NOTCH1*^+^), and one EVT state (*HLA-G*^+^, *ITGA1*^+^) were characterized (Fig. 1A; Fig. S1A, S1B).

**Figure 1:**
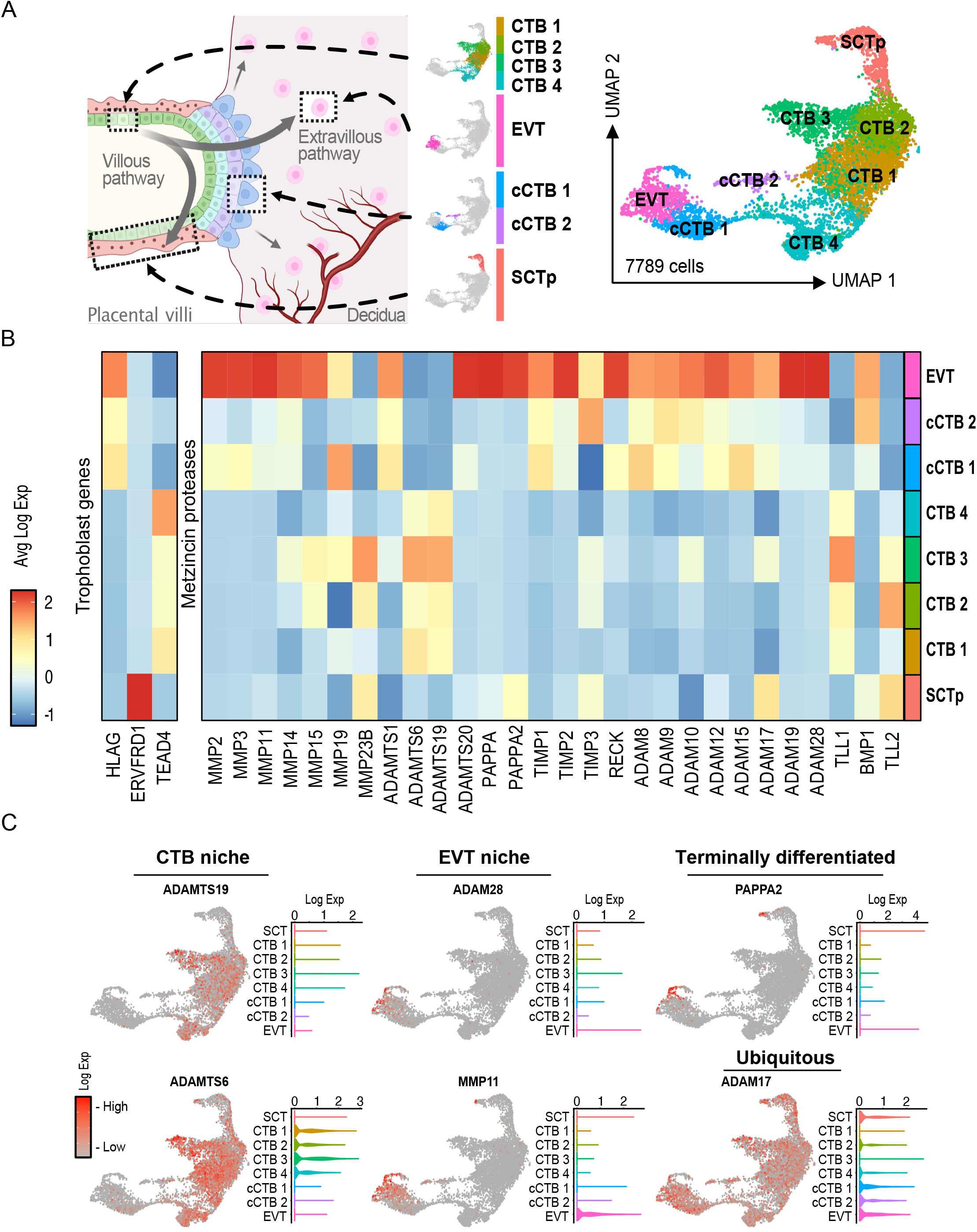
The protease landscape within the trophoblast interface. **(A)** Schematic of first trimester chorionic villous showing cytotrophoblast differentiation along the villous and extravillous pathways. Highlighted are trophoblast cell types: Cytotrophoblast (CTB), Extravillous trophoblast (EVT), Column cytotrophoblast (cCTB), Syncytiotrophoblast precursor (SCTp). Uniform Manifold Approximation Projection (UMAP) of 7789 captured trophoblasts including 8 trophoblast clusters: CTB1-4, EVT, cCTB1-2, SCTp. Approximate locations of *in silico-*denoted clusters are highlighted with black arrows. **(B)** Heatmap showing the average expression of 24 identified proteases in each trophoblast cluster. Trophoblast (sub)lineage identifying genes *HLA-G* (EVT), *TEAD4* (CTB), *ERVFRD-1* (SCT) are included. **(C)** Feature plots denoting cluster-associated gene expression and violin plots showing gene expression range of highly expressed proteases in CTB and EVT niches, in terminally differentiated cells, and in all clusters (ubiquitous).

To map metzincin protease gene signatures within these defined trophoblast states, a trophoblast state-based expression analysis of genes encoding human MMPs, astacins, ADAMs, ADAMTSs, and pappalysins, as well as inhibitors to these proteases, was performed (complete list of metzincin genes is shown in Table S1). Only metzincin genes expressed in ≥ 1% of cells across all 8 states or in ≥ 5% of cells within one specific state were selected for downstream analyses; this level of stringency accounts for the stochastic nature of scRNA-seq technologies and allows for the capture of proteases expressed in specific trophoblast sub-states as well as those expressed broadly across multiple states. Similar thresholds have been used elsewhere [30][29]. Using these parameters, we identified 24 metzincin proteases and 4 metalloprotease inhibitors in human first trimester trophoblasts with *TEAD4, ERVFRD1*, and *HLA-G* serving as reference marker genes for CTB, SCTp, and cCTB/EVT states, respectively (Fig. 1B). Specifically, cluster-averaged levels of metzincin genes show multiple MMPs (*MMP2*, -*3*, -*11*, -*14*, -*15*, -*19*, -*23B*), ADAMs (*ADAM8, -9, -10, -12, -15, -17, -19, -28*), ADAMTSs (*ADAMTS1, -6, -19, -20*), pappalysins (*PAPPA, PAPPA2*), and astacins (*TLL1, TLL2, BMP1*) to be expressed in first trimester trophoblasts (Fig. 1B). The four metalloprotease inhibitors *TIMP1, TIMP2, TIMP3*, and *RECK* are primarily confined to the EVT state and, to a lesser extent, the cCTB state (Fig. 1B). The majority of metzincin genes (19/24) are also preferentially expressed by invasive EVT (Fig. 1B; 1C). cCTB-1 and -2 states, developmentally upstream of EVT, likewise show broad expression of metzincin proteases, though levels are generally lower than those seen in EVT. *MMP19* is the exception by showing the highest expression levels in the cCTB1 state (Fig. 1B).

Though CTB progenitor states do not show as extensive expression of metzincin genes, a subset of MMPs (*MMP14, -15, -19, -23B*), ADAMs (*ADAM10, ADAM17*), ADAMTSs (*ADAMTS1, -6, -19*), and astacins (TLL1, TLL2) are expressed at moderate levels within the CTB1-4 states (Fig. 1B; 1C). Additionally, within the SCTp state, *MMP23B, PAPPA2, ADAM17, BMP1*, and *TLL2* are expressed by a substantial number of cells (Fig. 1B; 1C). Because of the broad ubiquitous expression of *ADAM17* across multiple trophoblast states (Fig. 1C), we examined the localization of ADAM17 by immunofluorescence microscopy (IF) in a separate cohort of first trimester placentas (Fig. 2A). Consistent with the scRNA-seq findings, ADAM17 signal was detected in EGFR^+^ CTB of floating chorionic villi and in HLA-G^+^ trophoblasts located proximally and distally, within placental anchoring columns (Fig. 2A). Due to non-specific antibody signal in the multinucleated hCG^+^ SCT (Fig. 2A; S2A), the strong immuno-labeling of ADAM17 within the outer SCT layer should be interpreted with caution. Nonetheless, quantification of the ADAM17 signal within the anchoring column showed that ADAM17 levels progressively increase along the EVT pathway, a finding consistent with our scRNA-seq analysis (Fig. 2B). ADAM17 signal was lowest in proximal anchoring column CTB and highest in HLA-G expressing distal column CTB (Fig. 2B). Overall, these findings capture the diversity of metzincin proteases expressed within trophoblast subtypes and distinct trophoblast states. Examples of proteases expressed within progenitor, in terminally differentiated EVT, and in SCT states are as well characterized at single cell resolution.

**Figure 2:**
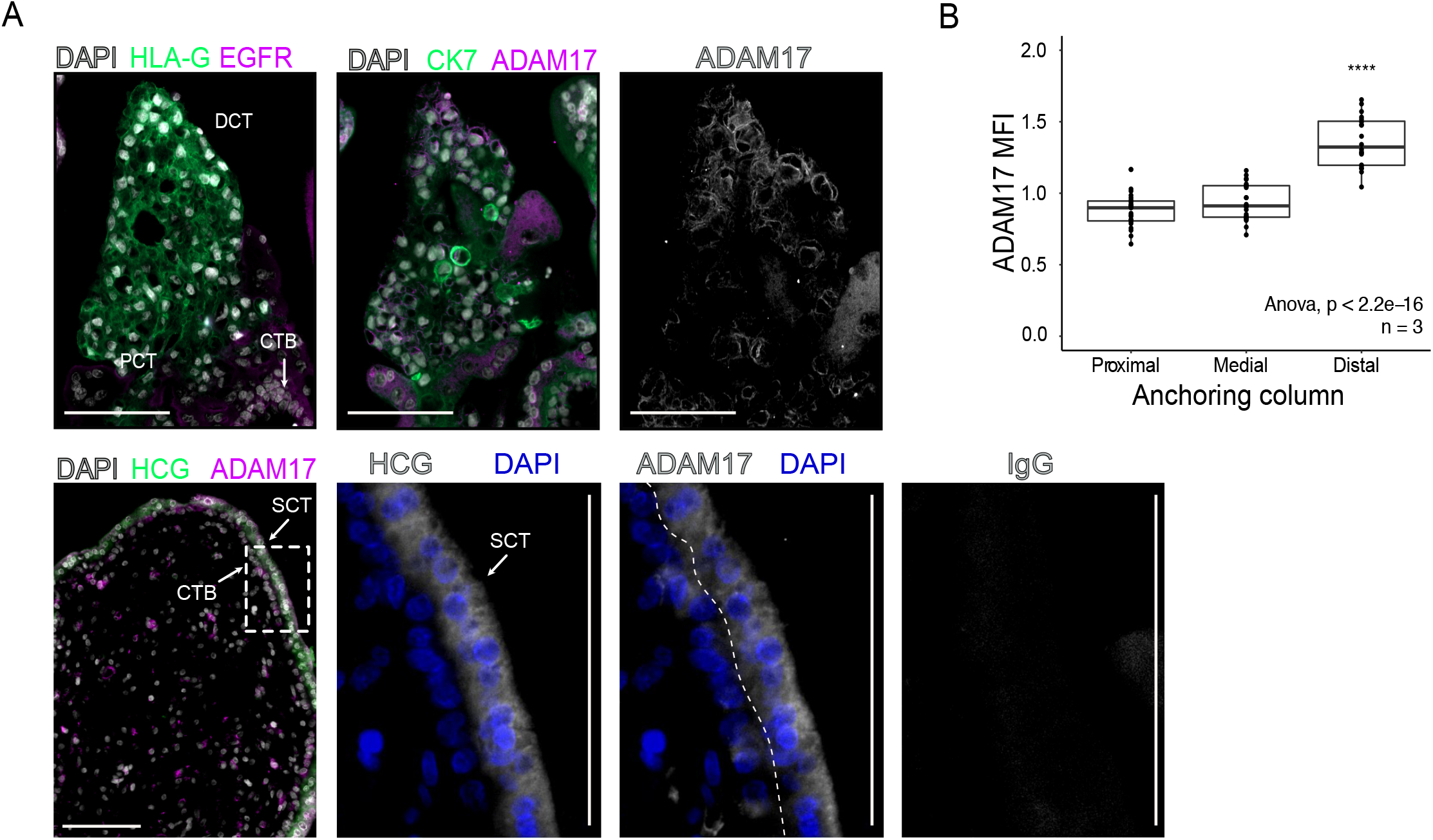
ADAM17 localization within first trimester chorionic villi. **(A)** Representative immunofluorescence (IF) images of HLA-G, EGFR, human chorionic gonadotrophin (HCG), cytokeratin 7 (CK7), and ADAM17 in first trimester placental anchoring villi. Villi labeled with polyclonal rabbit IgG in place of primary antibody serves as background signal control. Shown are proximal column trophoblasts (PCT), distal column trophoblasts (DCT), syncytiotrophoblasts (SCT), and cytotrophoblasts (CTB). Bar = 100 µm. **(B)** Mean fluorescent intensity (MFI) measurements of ADAM17 in proximal, medial, and distal first trimester anchoring placental villi section (n = 3). Statistical analyses between groups were performed using ANOVA and two-tailed Tukey post-test; significance *p* <0.05 **** = *p* <0.0001.

### Defining metzincin metalloprotease dynamics during trophoblast differentiation

Efforts to understand the roles of metalloproteases in trophoblast biology have primarily focussed on their involvements in promoting invasive and migratory EVT processes. However, given that metzincin proteases control multiple processes important in cell regeneration and fate specification (via modulation of substrate/receptor activity and availability, as well as cell-cell / cell-matrix adhesion), we next set out to examine how metzincin protease dynamics align with specific cell differentiation trajectories. Using the combined scRNA-seq datasets described above, we recapitulated Monocle 2 and Monocle 3 pseudotime cell trajectory modelling previously described in Shannon et al [26]. Monocle 2 semi-supervised ordering revealed two differentiation trajectories originating within the CTB1-4 states and developing towards mature EVT and SCTp states, respectively (Fig 3A; Fig. S2B). By sub-setting the 24 metzincin protease genes expressed in trophoblasts and aligning their expression levels along pseudotime-derived cell trajectories, we show protease expression kinetics during SCT and EVT differentiation (Fig. 3B). Plotting known stem/progenitor cell (*CDX2, TEAD4*), trophoblast column (*NOTCH1*), and terminally differentiated EVT (*HLA-G*) and SCT (*ERVFRD-1*) genes along pseudotime shows that Monocle 2 correctly reconstructs the anticipated cell state ordering during EVT and SCT differentiation (Fig. 3B). Specifically, cells expressing high levels of *CDX2* localize to the predicted pseudotime origin, whereas *TEAD4* expressing cells proximally flank this origin along both the EVT and SCT differentiation branches; as expected, *TEAD4* levels gradually decrease as cells mature into EVT and SCT (Fig. 3B). *NOTCH1*, a gene thought to play a central role in EVT progenitor renewal [42], shows the highest expression in trophoblasts found midway along the EVT branch. Further, *HLA-G* and *ERVFRD-1*, genes associated with more mature EVT and SCT states, respectively, show elevated expression in trophoblasts nearing EVT and SCT ordering endpoints (Fig. 3B). In summary, Monocle 2 pseudotime ordering recapitulates known transcriptomic events important in human trophoblast development.

**Figure 3:**
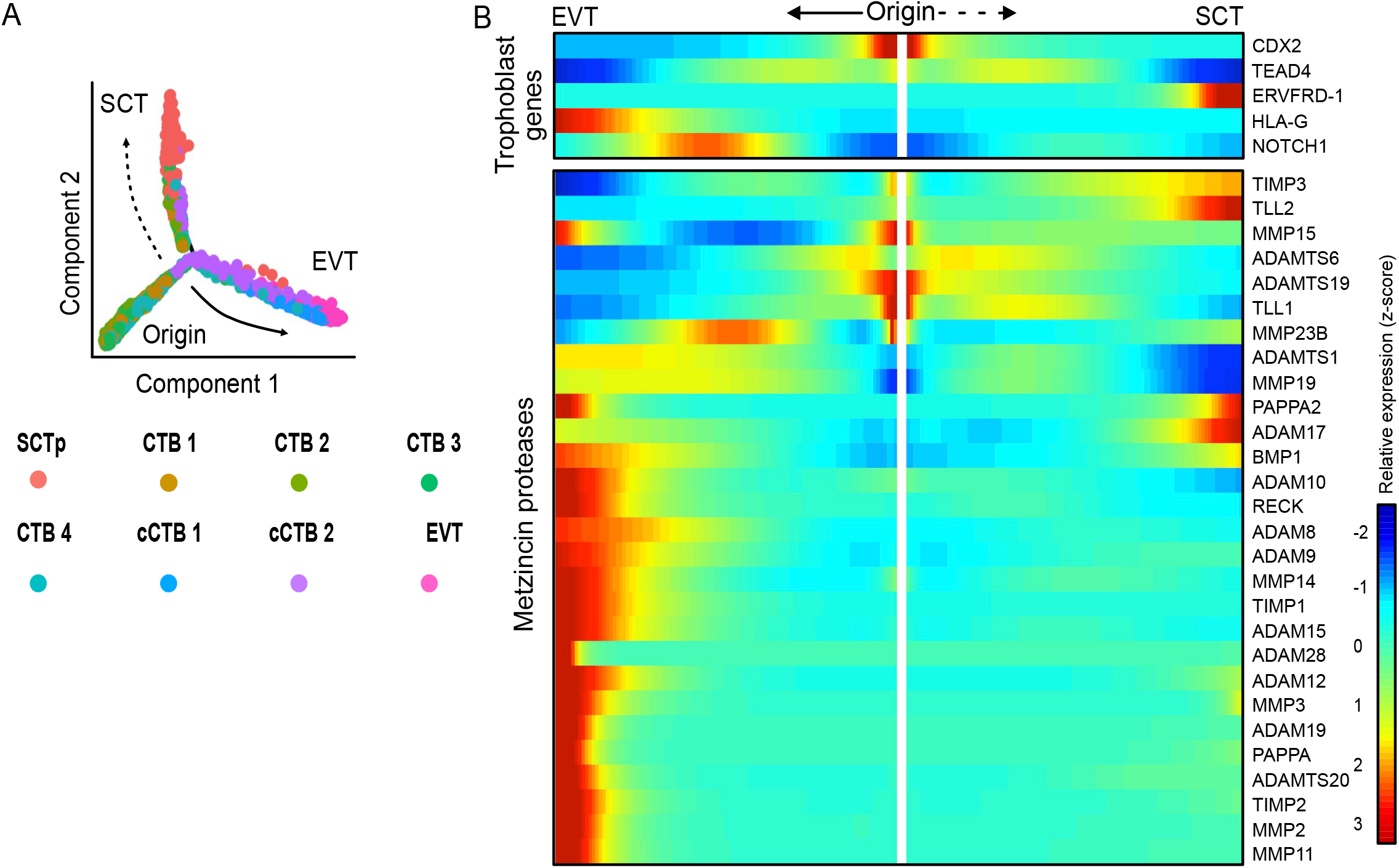
Metalloprotease dynamics along trophoblast lineage trajectories. **(A)** Monocle 2 pseudotime graph showing trophoblast cluster distribution along the villous and extravillous differentiation pathway. **(B)** Heatmap highlighting expression of 24 identified proteases and 4 protease inhibitors across Monocle2-informed pseudotime. Included within the heatmap are known genes that align with the CTB (*CDX2, TEAD4*), the SCT (*ERVFRD-1*), the cCTB (*NOTCH1*), and the EVT (*HLA-G*) states.

Alignment of the 24 metzincin proteases identified as expressed in trophoblasts along the pseudotime trajectory shows that expression of most genes (19/24) predominately skews towards the EVT state (Fig. 3B). This is consistent with previous descriptions of metzincin proteases like *MMP2* [19], *MMP14* [43], and *ADAM12* [20,21] in controlling EVT invasion. Notably, the expression of proteases like *ADAM8, ADAM9*, and *BMP1* appear upstream of *HLA-G*-expressing EVT, suggesting that these genes play roles in column trophoblast and immature EVT development (Fig. 3B). By contrast, the expression of other proteases (i.e., *ADAM28, PAPPA, MMP2, MMP15, ADAM19*) occurs much later along the EVT trajectory, suggesting that these genes may have roles specific to mature EVT functions like uterine stroma extracellular matrix degradation and invasion (Fig. 3B). As CTB states transition into SCT precursors, levels of *TLL2, PAPPA2*, and *ADAM17* markedly increase, indicating that these proteases play roles in aspects of SCT formation (Fig. 3B). Notably, *ADAM17, BMP1, MMP3*, and *PAPPA2* show bimodal-like patterns, where expression levels increase in cells transitioning along both the EVT and SCT pathways, suggesting that these proteases play complex roles in placental development (Fig. 3B).

Proximal to the defined pseudotime origin, a subset of proteases is observed to be expressed in primitive trophoblast states (Fig. 3B). *MMP15, MMP23B, ADAMTS19*, and *TLL1* all show high expression in cells closely aligning to the pseudotime origin and co-expressing the trophoblast stem cell/progenitor gene marker, *CDX2* (Fig. 3B). *ADAMTS6* levels align most tightly with cells directly flanking the origin, a pattern similarly observed for *TEAD4* (Fig. 3B). Examining the kinetics of metalloprotease inhibitor expression along EVT and SCT trajectories show that *RECK, TIMP1*, and *TIMP2* levels are highest in cells aligning with EVT endpoint states (Fig. 3B). *TIMP3* levels, on the other hand, are highest at the origin and in cell states gradually progressing along the SCT trajectory (Fig. 3B). Together, this data highlights metzincin protease and inhibitor gene expression dynamics in human trophoblasts differentiating along the SCT and EVT pathways. These findings provide insight into possible functions and the developmental timing of specific proteases expressed across trophoblast regeneration and differentiation.

### Progenitor metzincin proteases and their kinetics within the CTB niche

Progenitor CTB must undergo continual renewal and, when appropriate, commit to differentiation along either the SCT or EVT pathways. Within this CTB niche, little is known about the role metzincin proteases play. However, the scRNA-seq analyses described above identified 11 metzincin encoding genes expressed in the CTB1-4 states. To examine the kinetics of CTB-associated protease expression more closely, we applied RNA velocity (scVelo) [35], an algorithm that measures relative abundances of unspliced versus spliced mRNA transcripts to predict cellular state progression. In line with scVelo modeling performed previously on this dataset [26], a predicted origin overlapping with the CTB2 state was identified with multiple paths extending from this origin towards the CTB3, SCTp, and EVT states (Fig. 4A). Assessment of the key progenitor/stem cell-associated genes *TEAD4* and *CDX2* within UMAP projections shows their expression to be largely defined within CTB progenitor states, though *TEAD4* expression is most frequent within the proliferative *MKI67*^+^ CTB4 state (Fig. 4B). Visualization of metzincin proteases identifies three broad expression patterns: CTB-specific, ubiquitous, and bimodal/transient (Fig. 4C-E). Metzincins predominantly expressed in CTB progenitor states are *ADAMTS6, ADAMTS19, MMP23B*, and *TLL1*, where *ADAMTS6* and *ADAMTS19* are expressed at higher levels, and in more cells per CTB state, than *MMP23B* and *TLL1* (Fig. 4C). Proteases expressed ubiquitously across all states, including CTB, are *ADAM10, ADAM17, ADAMTS1, MMP14*, and *MMP19* (Fig. 4D). However, it is notable that while expression is observed in all trophoblast states, levels of *ADAM10, ADAMTS1, MMP14*, and *MMP19* are markedly higher in cells aligning with column CTB (cCTB1, cCTB2) and EVT states (Fig. 4D). Lastly, *TLL2* and *MMP15* show transient levels of expression as trophoblasts differentiate along the EVT and SCT pathways. *TLL2* levels are prevalent in trophoblasts aligning to the CTB origin and show elevation in the differentiated SCTp state (Fig. 4E). *MMP15* also shows a high frequency of expression in origin-associated trophoblast states but this differs from *TLL2* with *MMP15* being more highly expressed in terminally differentiated EVT (Fig. 4E). Together, this data identifies specific metzincin protease genes expressed within CTB states and illustrates how levels of these genes change within unique CTB states and along SCT and EVT pathways.

**Figure 4:**
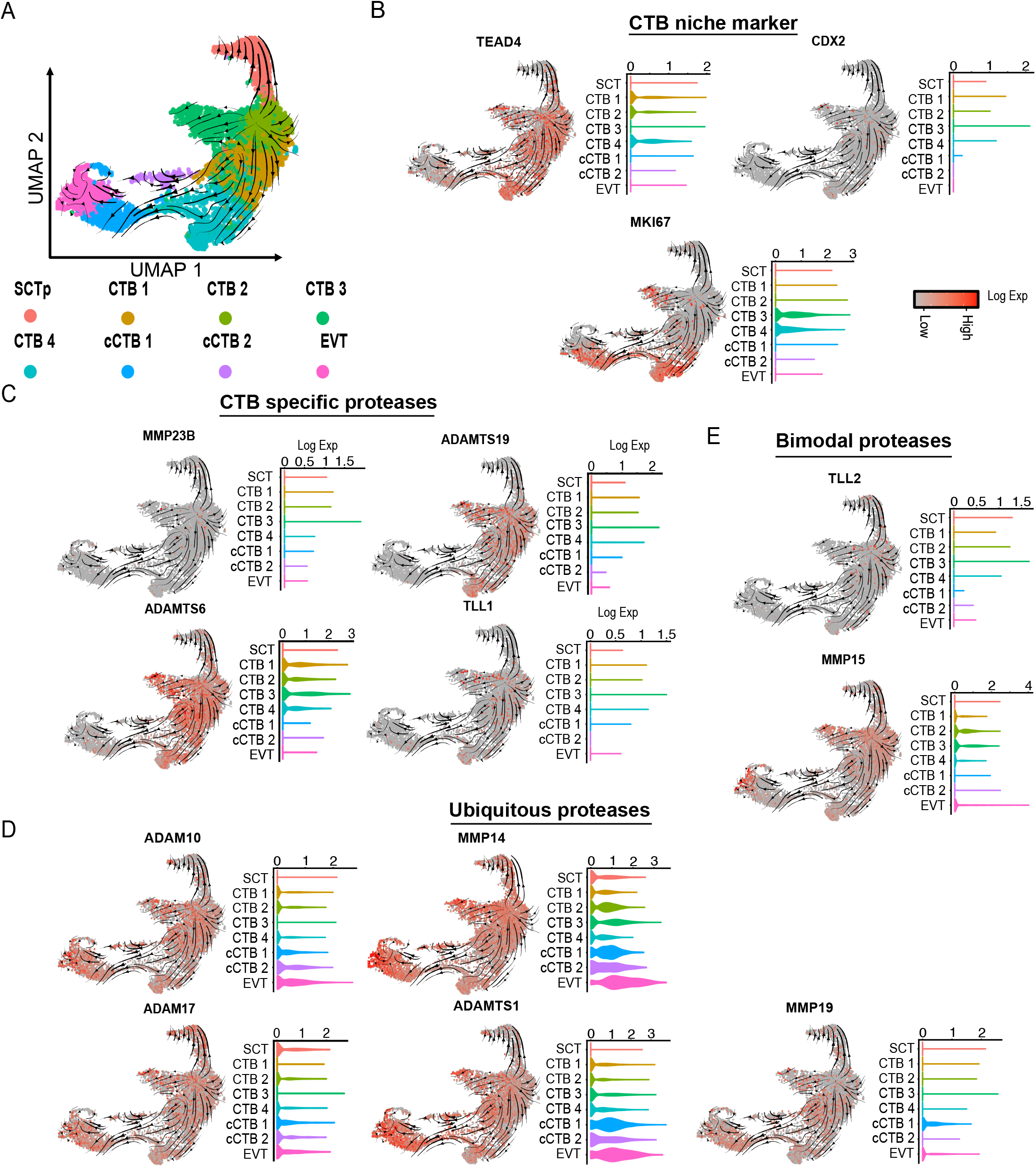
Metzincin proteases and the CTB niche. **(A)** Feature plot showing trophoblast cell states overlain with the generated RNA velocity vector field (arrows), designating trophoblast differentiation directionality. Feature plots denoting cluster specificity and violin plots showing gene expression range of **(B)** prominent CTB niche markers, **(C)** CTB specific proteases, **(D)** ubiquitous proteases, and **(E)** bimodal proteases overlain with the generated RNA velocity vector field (arrows), designating trophoblast differentiation directionality.

### Mapping the metzincin-substrate landscape within the CTB niche

Our finding that multiple metzincin proteases are preferentially expressed within progenitor CTB sheds light into the possible roles these metalloproteases play in trophoblast progenitor renewal and the early steps of trophoblast differentiation. To investigate the biological functions controlled by metzincins within the CTB niche, we set out to examine enzyme-substrate interactions. Focussing on only metzincin genes expressed in ≥ 20% of CTB (so as to improve interpretability of protease-substrate relationships, as performed in other studies [44]); we restricted our analyses to *ADAM10, ADAM17, ADAMTS6, ADAMTS1, MMP14*, and *MMP15*. We used the NicheNet package which combines dataset-specific gene expression data with existing knowledge on ligand-receptor, signaling, and gene regulatory networks. Since our analysis focussed solely on direct protease interactions with substrates and other binding partners, we excluded data sources containing downstream gene regulatory information, indirect interactions, phosphorylation events, and kinase-substrate interactions (Table S2). We further manually added verified protease-substrate interactions specific to *ADAM10, ADAM17, ADAMTS6, ADAMTS1, MMP14*, and *MMP15* in response to the non-exhaustive nature of the NicheNet curated directory (Table S3).

From the protease-substrate interaction list, only genes encoding substrates expressed in ≥ 20% of CTB were selected. In total, 154 potential protease-substrate pairs are identified in the CTB niche (Fig. 5A) (Table S3). Not surprising, substrates common to both *ADAM10* and *ADAM17* are identified, including *ITGB1, CDH1, APP* and *EGFR* (Fig. 5A). Interaction potential scoring highlights *PTPN3* (tyrosine phosphatase), *TNFRSF1A*, and *MAD2L1* as the substrates most strongly interacting with *ADAM17* (Fig. 5A). *ADAM10*, on the other hand, shows interaction with *MET, DLG1, ITM2B*, and *B3GNT2* within the CTB1-4 states (Fig. 5A). *MMP14* and *MMP15* show only moderate overlap in substrates such as *TGM2, LAMB1*, and *LAMC1*, where *MMP14* itself is suggested to have 75 different substrates and interactions (Fig. 5A). Both *ADAMTS1* and *ADAMTS6* only have 7 and 11 substrates or interactions in the CTB niche, respectively (Fig. 5A). This may be partly because *ADAMTS6* had been designated an orphan protease until very recently [45]. Notably, both *ADAMTS1* and *ADAMTS6* interact with substrates that encode proteins important for elastic fiber assembly, such as *MFAP5, FBLN1, FBN2, FBN1*, and *SDC4* (Fig. 5A). Gene Ontology analysis on the 154 genes encoding CTB metzincin substrates highlights broad functional roles in regulating focal adhesions and cell substrate junction organization (Fig. 5B); the latter referring to specialized regions of connection between a cell and the extracellular matrix. These top ontology hits suggest that CTB progenitor metzincins control the turnover of the extracellular environment.

**Figure 5:**
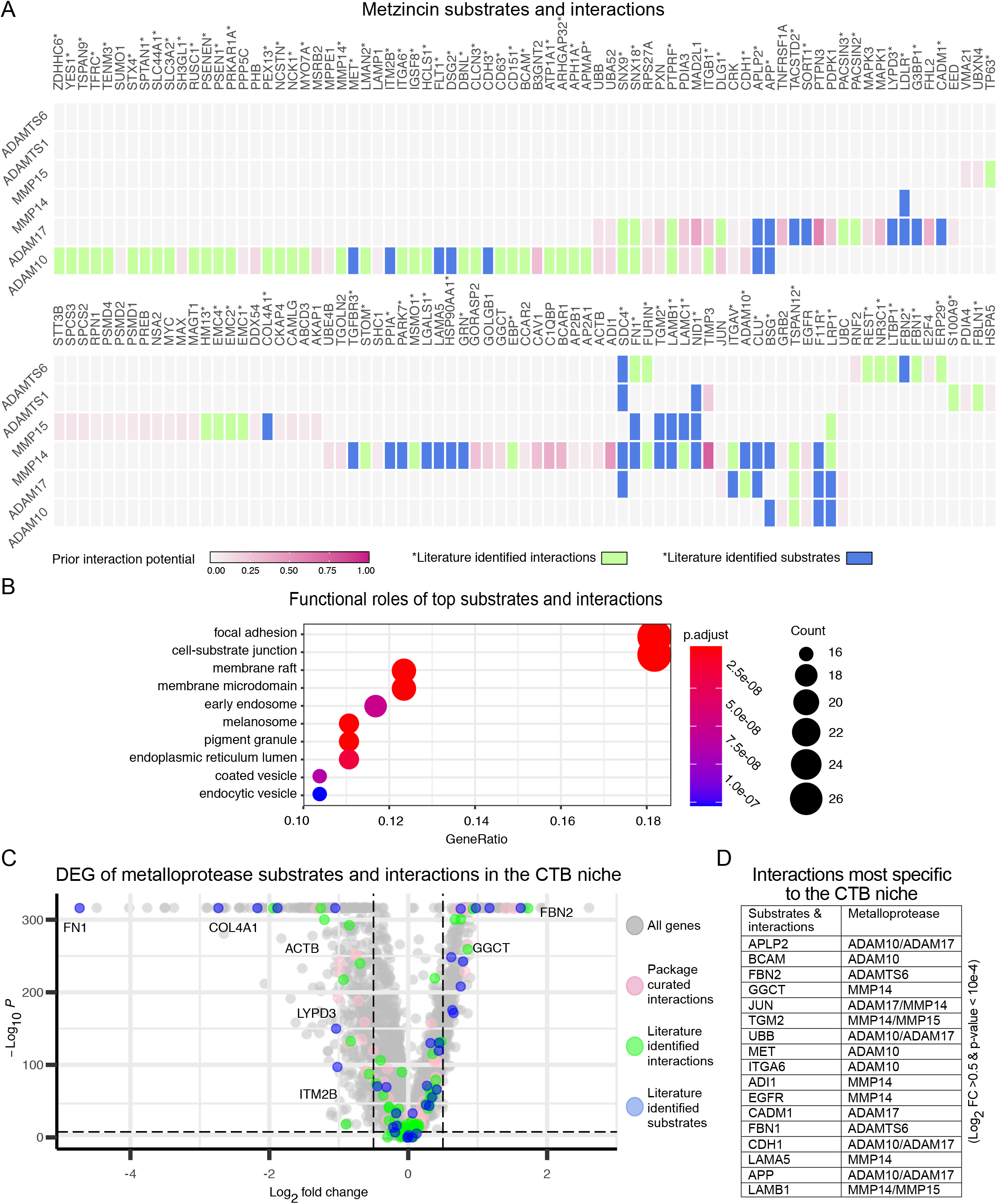
CTB metalloprotease substrates and downstream targets. **(A)** Heatmap showing NicheNet-identified potential interactions between CTB proteases and their respective substrates and binding partners in CTB cluster 1-4. Package curated interaction potentials are marked with pink colour intensity. Literature identified interactions (which possess no prior interaction potential) are marked in lime green. Literature identified substrates are marked in blue. **(B)** Gene ontology results showing pathways enriched for identified substrates and interactions (*n*=154 genes). **(C)** Volcano plot showing differentially expressed genes between CTB1-4 and other trophoblast clusters (cCTB 1-2, EVT, SCTp). Previously identified substrates, bindings partners and indirect targets are highlighted in pink (NicheNet-curated), lime green (library curated interacting partners) or blue (library curated substrates). **(D)** Table showing top identified substrates and interactions (avg log fold change > 0.5 & p value < 10e-4) and their corresponding proteases.

To focus on highly expressed metzincin substrates showing preferential expression in the CTB niche, we next performed a differential gene expression (DEG) analysis comparing mRNA levels of the 154 NicheNet-identified substrates and binding partners within CTB 1-4 to the other four non-CTB trophoblast states (i.e., cCTB1, cCTB2, EVT, SCT) (Fig. 5C). DEG analysis identifies 17 genes encoding 11 metzincin substrates and 6 physical interaction partners enriched within the CTB niche (Fig. 5D). CTB-specific protease-substrate pairs are summarized in tabular format, where the top three most highly expressed substrates or binding partners, ordered by fold change (FC) and *p* value ranking, are *APLP2* (encoding amyloid-like protein 2, a target of ADAM10 and ADAM17), *BCAM* (encoding basal cell adhesion molecule, a target of ADAM10), and *FBN2* (encoding fibrillin-2, a target of ADAMTS6) (Fig. 5D). Overall, expression of 154 genes encoding prominent substrates or binding partners of highly expressed CTB proteases was identified. This interaction matrix provides insights into possible roles proteases may play in the CTB niche.

### *ADAMTS6* and *FBN2* are co-expressed by CTB progenitors

*FBN2* was the third most highly expressed metzincin protease substrate specific to the CTB niche. As indicated above, *FBN2* encodes for the protein fibrillin-2, a large extracellular matrix protein important in the formation of microfibril filaments that serve as scaffolds for other structural proteins [46]. Importantly, fibrillin-2 is proteolytically degraded by ADAMTS6 [45], a protease whose expression also aligned preferentially to the CTB niche. Therefore, we set out to verify that both *ADAMTS6* and *FBN2* are specific and co-expressed within CTB progenitors of the human first trimester placenta. When examining expression of *ADAMTS6* and *FBN2* in CTB1-4 states (comprising 5477 cells), 2301 cells show co-expression of both genes (Fig. 6A). Only 159 CTB solely express *ADAMTS6*, whereas approximately 50% of CTB (2654 cells) express *FBN2* independent of *ADAMTS6* (Fig. 6A). Importantly, both *ADAMTS6* and *FBN2* show similar expression patterns within the eight scRNA-seq defined trophoblast states, where highest expression levels of either gene are found in CTB3 (Fig. 6B).

**Figure 6:**
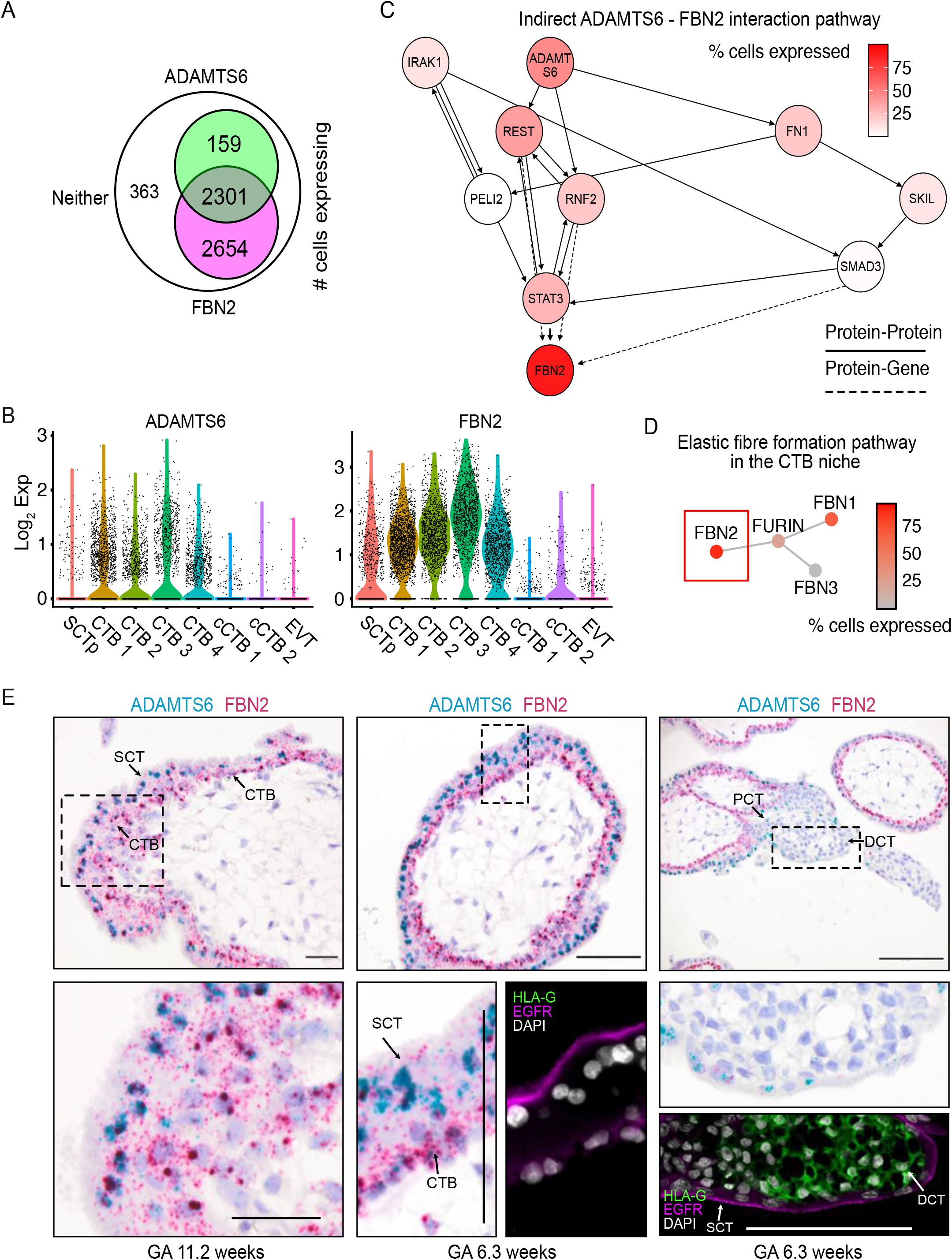
ADAMTS6 and FBN2 characterization within the CTB niche. **(A)** Volcano plot highlighting *FBN2* specificity to the CTB niche compared to all other differentially expressed genes between CTB1-4 and other trophoblast clusters (cCTB 1-2, EVT, SCTp). **(B)** Diagram illustrating cells expressing neither *ADAMTS6* nor *FBN2* (white), *ADAMTS6* expressed alone (green), *FBN2* expressed alone (pink), or *ADAMTS6* and *FBN2* co-expression (grey). **(C)** Violin plot showing expression of *ADAMTS6* and *FBN2* within identified trophoblasts. Cluster identities are indicated by colour. **(D)** Diagram illustrating alternative interaction pathways between ADAMTS6 and FBN2 as calculated by nichenet. Percentage of CTB cells expressing relevant genes are marked in red. Solid arrows designate directionality of protein-protein interactions. Dashed arrows designate directionality of protein-gene interactions. **(E)** Percentage of CTB cells expressing genes interacting with FBN2 in the elastic fibre formation pathway (Reactome pathway database). **(F)** Representative *in situ* hybridization images showing mRNA expression of *ADAMTS6* and *FBN2* in floating and anchoring villi of early (6.3 weeks) and late (11.2 weeks) first trimester chorionic villi. Proximal column trophoblasts (PCT), distal column trophoblasts (DCT), cytotrophoblasts (CTB) and syncytiotrophoblasts (SCT) are indicated. Serially sectioned villi labeled with HLA-G and EGFR help identify column and cytotrophoblast compartments; nuclei are labeled with DAPI. Bar = 100 µm.

Though fibrillin-2 is a direct substrate of ADAMTS6, it is also possible that indirect interactions between ADAMTS6 and fibrillin-2 may exist. To assess this, we examined the shortest ligand-target signalling path (protease-target in this case) using NicheNet (all NicheNet-curated data sources were included). We found that *ADAMTS6* and *FBN2* may potentially interact indirectly through *REST, RNF2*, and *STAT3*, as these genes were moderately expressed in CTB progenitors (Fig. 6C). NicheNet analysis further indicated gene signalling interactions between *PELI2, SMAD3, SKIL, FN1*, and *IRAK1*, stemming from *ADAMTS6*-*FN1* interaction and ADAMTS6-related control of the transcription factor REST (Fig. 6C). However, the potential for an interaction between *ADAMTS6* and *FBN2* via *FN1* is unlikely due to the low expression of *SMAD3* in all four CTB states (Fig. 6C). Because fibrillin-2 is involved in microfibril/elastic fiber formation and related tissue resiliency, we next examined the relationship and expression of other genes central to elastic fiber formation within the CTB niche (Fig. 6D). We focussed on members directly interacting with *FBN2* (Fig. 6D) and indirectly interacting with *FBN2* (Fig. S3). The majority of microfibril-linked genes are expressed in CTB states at high levels (e.g., *FBN1, EFEMP1, FBLN1, MFAP1*) (Fig. 6D), suggesting that elastic fibers/microfibrils play a role in CTB extracellular matrix scaffolding and the possible establishment or maintenance of the CTB niche.

Lastly, we set out to verify *ADAMTS6* and *FBN2* expression and localization within *in vivo* chorionic villi of first trimester placenta. *In situ* hybridization of *ADAMTS6* and *FBN2* in early (6.3 weeks’ gestation) and late (11.2 weeks’ gestation) placentas show that both genes are highly expressed within the different cell types of floating villi (Fig. 6E). Consistent with scRNA-seq findings, *FBN2* shows a robust signal within EGFR^+^ progenitor CTB from a serial section of the same placenta; little/no *FBN2* signal is detected within HLA-G^+^ anchoring column trophoblasts (Fig. 6E). However, *FBN2* is expressed, albeit at a lower level, within the multinucleated syncytiotrophoblast (Fig. 6E). The spatial distribution of *ADAMTS6* within chorionic villi is somewhat surprising. Consistent with scRNA-seq data, moderate signal of *ADAMTS6* is observed in CTB progenitors, and low/no *ADAMTS6* signal is seen within cells of the anchoring columns (Fig. 6E). However, high levels of *ADAMTS6* are shown within the syncytiotrophoblast (Fig. 6E), likely reflecting a limitation of scRNA-seq in capturing multinucleated structures and cells. Together, this work highlights the expression of *ADAMTS6* and *FBN2*, as well as their interacting genes and processes, in the first trimester human placenta. Further, these findings provide evidence that the ADAMTS6-fibrillin-2 protease-substrate pair is co-expressed and largely specific to trophoblasts residing within the progenitor CTB niche.

## DISCUSSION

Using established scRNA-seq datasets of first trimester chorionic villi, this work characterizes the metzincin protease landscape in diverse subtypes and states of human trophoblast. Twenty-four metzincin proteases and four metalloproteinase inhibitors were mapped to distinct trophoblast subtypes. State-of-the-art single cell lineage trajectory analyses then aligned metzincin expression dynamics along both the villous and extravillous differentiation pathways, as well as within the CTB progenitor niche. While metzincin protease expression showed a strong expression bias towards cells of the EVT lineage, our analyses identified 11 metalloproteases expressed by trophoblasts overlapping with CTB progenitor states. Within CTB progenitors, we provide a predicted framework of possible metzincin-substrate interactions, highlighting the importance of elastic fiber/microfibril remodeling processes within the CTB progenitor niche. Specifically, we show co-expression of *ADAMTS6* and *FBN2*, a recently identified protease-substrate pair, in CTB. Together, this work comprehensively characterizes the metzincin landscape in human trophoblasts and provides an important resource for future studies investigating the roles of metzincin proteases in placental development and function.

Previous studies have explored the roles of metzincin proteases in trophoblast biology and have spatially defined MMP2 [19,47], MMP3 [48], MMP14 [49], MMP15 [50], ADAM8 [51], ADAM12 [20–22], ADAM28 [52], PAPPA [53], and PAPPA2 [54] to subtypes of trophoblast within first trimester chorionic villi. Single cell transcriptomic assessment of these aforementioned metzincins performed in this study overall supports the conclusions of their previous spatial characterizations, with the majority of these proteases localizing to column trophoblasts and invasive EVT. Absent from this list is MMP9, a matrix metalloprotease often described as an EVT marker important in controlling trophoblast invasion [55]. It should be noted that the majority of (if not all) studies describing MMP9 expression specificity to invasive trophoblast subsets characterized and tested MMP9 function in immortalized trophoblastic cell lines [56] or inadequately described primary trophoblast cultures [57]. This latter example is especially important as primary cultures of CTB and EVT are challenging to establish and experiment on. In part, this is because primary cultures do not passage or proliferate well in culture, and as a result of this, are often quickly over-run by contaminating placental fibroblasts [3]. Therefore, it is essential for studies aiming to examine the importance of specific genes or pathways in trophoblast processes to carefully confirm trophoblast identity using recently accepted molecular and genetic trophoblast-identifying characteristics [58] when deriving primary cultures.

Of the 24 metzincin proteases ascribed to various trophoblast states, to our knowledge, 15 have not been identified in first trimester trophoblasts, and include MMPs (*MMP11, -19, -23B*), ADAMs (*ADAM9, - 10, -15, -17, -19*), ADAMTSs (*ADAMTS1, -6, -19, -20*), and astacins (*TLL1, TLL2, BMP1*). Notably, PAPPA and ADAM17 have been localized to the syncytiotrophoblast of term placentas, where their expression is thought to control growth factor availability [59,60] and disruptions in their regulation may contribute to the development of preeclampsia [59,61]. Verification of ADAM17 localization by immunofluorescence imaging supports previous findings that ADAM17 localizes to the syncytiotrophoblast. However, our findings further elaborate on the possible functions of ADAM17 in the placenta in that its expression is also detected in progenitor CTB, anchoring column trophoblast, and invasive EVT. Notably, *ADAM17* levels increase in trophoblasts as they progress along both the villous and extravillous pathways, suggesting that it may coordinate complex processes linked to cell migration and cell-cell fusion. Given the diverse substrates of ADAM17, both in the context of health and disease, it will be challenging to carefully assign specific biological functions to ADAM17 in trophoblast development and function. Similar to ADAM17, other metzincins show a bimodal-like pattern of expression as trophoblast differentiation terminates along the villous and extravillous pathways; *PAPPA2* and *BMP1* are both highly expressed in EVT and syncytiotrophoblast precursors. BMP1 controls fibrillar collagen assembly by cleaving C-propeptides of procollagens I-III and also regulates the availability of TGFβ-like proteins that are critical in tissue patterning during development [62]. PAPPA2 on the other hand likely regulates IGF-1 and IGF-2 availability and latency through cleavage of IGFBP5 [63] and in turn this cleavage may control trophoblast proliferation and survival, as well as coordinate placental organ growth.

Recent findings have highlighted the importance of metzincin proteases in the mesenchymal [64], hematopoietic [65] and skeletal muscle [66] stem cell niche. Knowledge on the actions of metzincin proteases in the cytotrophoblast stem cell niche are, to our understanding, nonexistent. CTB progenitors are sandwiched between a laminin-rich basal lamina and an overlying syncytium monolayer. This spatial arrangement limits the source of soluble factors that interact with CTB. Most likely, soluble factors originate from the overlying syncytiotrophoblast or from within the CTB niche itself. In this study we identify 11 metzincins that are expressed by CTB, though only *MMP15, MMP23B, ADAMTS6, ADAMTS19*, and *TLL1* are specifically aligned within CTB. Expression of *MMP15, MMP23B, ADAMTS19*, and *TLL1* align with the Monocle2 predicted CTB origin, while *ADAMTS6* expression is most specific to the CTB3 progenitor state, suggesting that these proteases likely regulate processes central to trophoblast stem cell maintenance. The concept of both membrane-bound and secreted proteases as important factors in controlling stem cell regeneration and maintenance has been gaining traction in recent years. For example, smooth muscle cell produced MMP17 controls intestinal stem cell proliferation through cleavage of matricellular proteins like periostin [67]; while ADAMTS18 regulates the mammary myoepithelial stem cell niche through the progesterone-Wnt4 controlled processing of collagen [23].

Our finding that ADAMTS6 and its recently identified substrate, fibrillin-2 [45], are highly and specifically expressed in CTB states suggests that ADAMTS6 may also play a role in regulating CTB progenitor maintenance. Fibrillin-2 is a major component of microfibril elastic fibers that are components of the extracellular matrix and confer elasticity and resilience to a tissue [68]. Importantly, morphogens like TGFβ and BMP family members are sequestered within microfibril elastin matrices, where proteolytic cleavage of microfibrils liberates these growth factors and in turn generates growth factor gradients important in tissue patterning and cellular development [69]. It is tempting to speculate that ADAMTS6 may be coordinating similar events within the CTB niche. Future research will need to focus on the importance of ADAMTS6 in CTB progenitor biology, where newly developed regenerative trophoblast platforms like human trophoblast stem cell lines [70] and organoids [58,71] can help in defining the interplay between ADAMT6 and fibrillin-2.

In addition to the ADAMTS6-fibrillin-2 interaction within the CTB niche, this study also highlights 154 substrates expressed in CTB that pair with CTB-expressed metzincin proteases. The high selection threshold applied in our NicheNet modeling ensured inclusion of probable and highly influential interactions. GO analysis of these 154 substrates identified focal adhesion and cell-matrix interaction pathways to be the most dominant and represented, suggesting that protease-substrate interactions in the progenitor trophoblast niche are impacting cell-matrix adhesion, integrin signaling, and matrix-controlled cell polarity. Interestingly, *BCAM*, encoding a cell adhesion molecule recently identified as being enriched in progenitor CTB in the first trimester [26], was shown in our analyses to be a top binding partner expressed preferentially in CTB and found to be specifically interacting with ADAM10. BCAM enables cell-basal lamina interactions by binding to laminin α5 chains of heterotrimeric laminin, a major constituent of the basement membrane ECM [72]. A recent study implicated laminin-1 as one of the key drivers of cell fate switching in human trophoblast stem cells (hTSC), where addition of laminin-1 to the culture media switched the directionality of progenitor trophoblast differentiation from the villous pathway towards the extravillous pathway [73]. Our dataset specifically identifies the cleavage of *LAMA5* and *LAMB1* by *MMP14* and *MMP15* within the cytotrophoblast niche, further highlighting laminin as a driver of trophoblast stem cell maintenance. This emphasizes the dependency of trophoblast cell fate decisions on extracellular cues. Holistically, our current study will function as an important resource in identifying additional metzincin-substrate interactions that are possibly important in CTB function.

An important caveat of the single-cell RNA sequencing approach is created by the current inability to sequence multinucleated cells contained in the syncytium. As exemplified by *ADAMTS6*, scRNA seq was unable to show the enrichment of this protease in the syncytium. This led us to miss important insights about SCT differentiation dynamics. Future single-nuclear sequencing approaches may overcome this limitation. The stochastic nature of gene expression paired with the high dropout rate that is commonly observed in single cell sequencing decreases the reliability of some of the findings made, especially with respect to genes expressed at comparatively low levels. By this reasoning, it is probable that previously described metalloproteases such as MMP9 do play roles in trophoblast biology albeit not captured by our current analysis.

Here we put in place a comprehensive resource for the investigation of metzincin proteases in first trimester human trophoblasts. We have effectively characterized the first trimester metzincin protease landscape by investigating metzincin expression in transcriptomically defined trophoblast states, aligned trophoblast-expressed metalloproteases to specific lineage trajectories, and identified known protease-substrate interactions that are likely taking place within the CTB progenitor niche. As a valuable resource to developmental and placental biologists, we hope that future work will be able to verify and test metzincin– substrate interactions to further elucidate the influence of ECM constituents in regulating trophoblast stemness and regeneration.

## Abbreviations

CTB: Cytotrophoblast
DCT: Distal column trophoblast
DGE: Differential gene expression
EVT: Extravillous trophoblast
GO: Gene ontology
HLA-G: Human leukocyte antigen G
IF: Immunofluorescence
scRNA-seq: single cell RNA sequencing
SCT: Syncytiotrophoblast
UMAP: Uniform manifold approximation and projection
ADAM: A disintegrin and metalloproteinase
MMP: Matrix metalloprotease
ADAMTS: A disintegrin and metalloproteinase with thrombospondin motifs
ECM: Extracellular cell matrix
EGFR: Epidermal growth factor receptor

## DATA AVAILABILITY/ACCESSION NUMBER

The ArrayExpress accession number for the publicly available data reported in this paper is: E-MTAB-6701. The GEO accession number for publicly available data reported in this paper is: GSE174481.

## AUTHOR CONTRIBUTIONS

AGB and JW conceptualized the study. AGB and JW designed the research. JW, BC, JB performed experiments and analysed data. JW and MJS prepared computational datasets and bioinformatics pipelines. AGB and JW wrote the paper. All authors read and approved the manuscript.

## ACKNOWLEDGEMENTS

The authors extend their sincere gratitude to the hard work of staff at British Columbia’s Women’s Hospital’s CARE Program for recruiting participants to our study. We also wish to acknowledge Jenna Treissman, Desmond Hui, and Dr. Hoa Le for their contribution to the establishment of the 10x single-cell RNA seq datasets.

## SUPPLEMENTAL FIGURE LEGENDS

**Supplemental Figure 1:**

(A) Uniform Manifold Approximation Projection (UMAP) of 7789 captured trophoblasts incl. 8 trophoblast clusters: CTB1-4, EVT, cCTB1-2, SCTp. (B) Feature plots denoting gene expression cluster specificity of key trophoblast markers.

**Supplemental Figure 2:**

(A) Representative IF image showing polyclonal rabbit IgG signal in chorionic anchoring and floating villi. Shown are proximal column trophoblasts (PCT), distal column trophoblasts (DCT), cytotrophoblasts (CTB) and syncytiotrophoblasts (SCT). Bar = 100 µm. (B) UMAP plot with monocle 3 trajectory graph overlain. Relative pseudotime values are color coordinated.

**Supplemental Figure 3:**

(A) Relative expression of 11 identified CTB proteases across pseudotime towards either the mature EVT (solid line) or SCT states (dashed line). Corresponding cell clusters are marked by colour. Trophoblast lineage specific genes *HLA-G* (EVT), *TEAD4* (CTB), and *ERVFRD-1* (SCT) are included.

**Supplemental Figure 4:**

(A) “Elastic fibre formation” pathway colored with gradient (red) indicating proportion (%) of cells expressing specific genes in CTB clusters 1-4.

