## Supplementary figures and images for "Transcriptomic mapping of the metzincin landscape in human trophoblasts"

### Supplemental Figure 1

#### Supplemental Figure 1

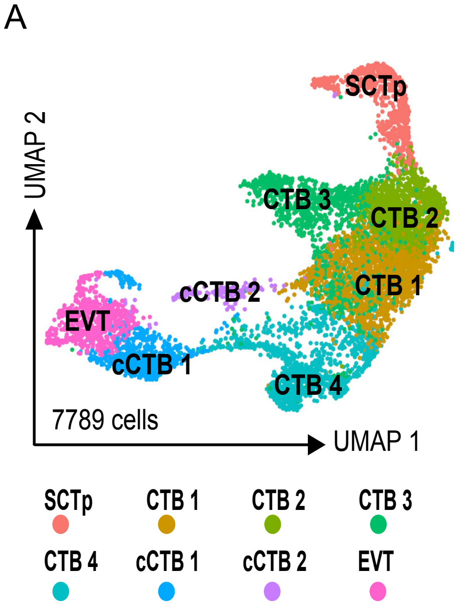


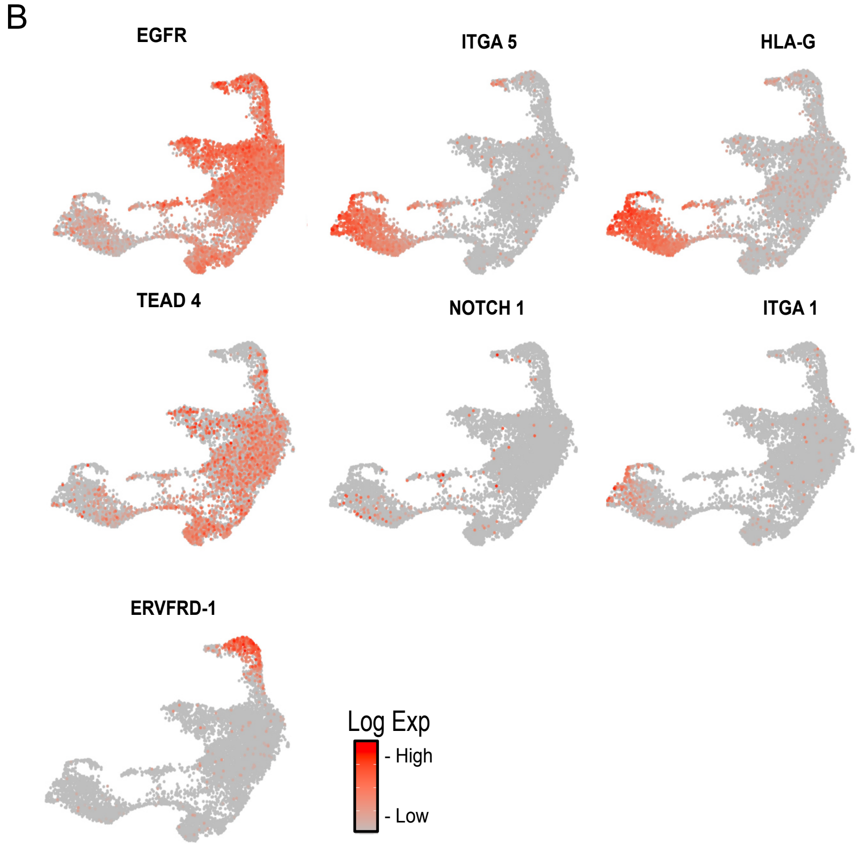

### Supplemental Figure 2

#### Supplemental Figure 2

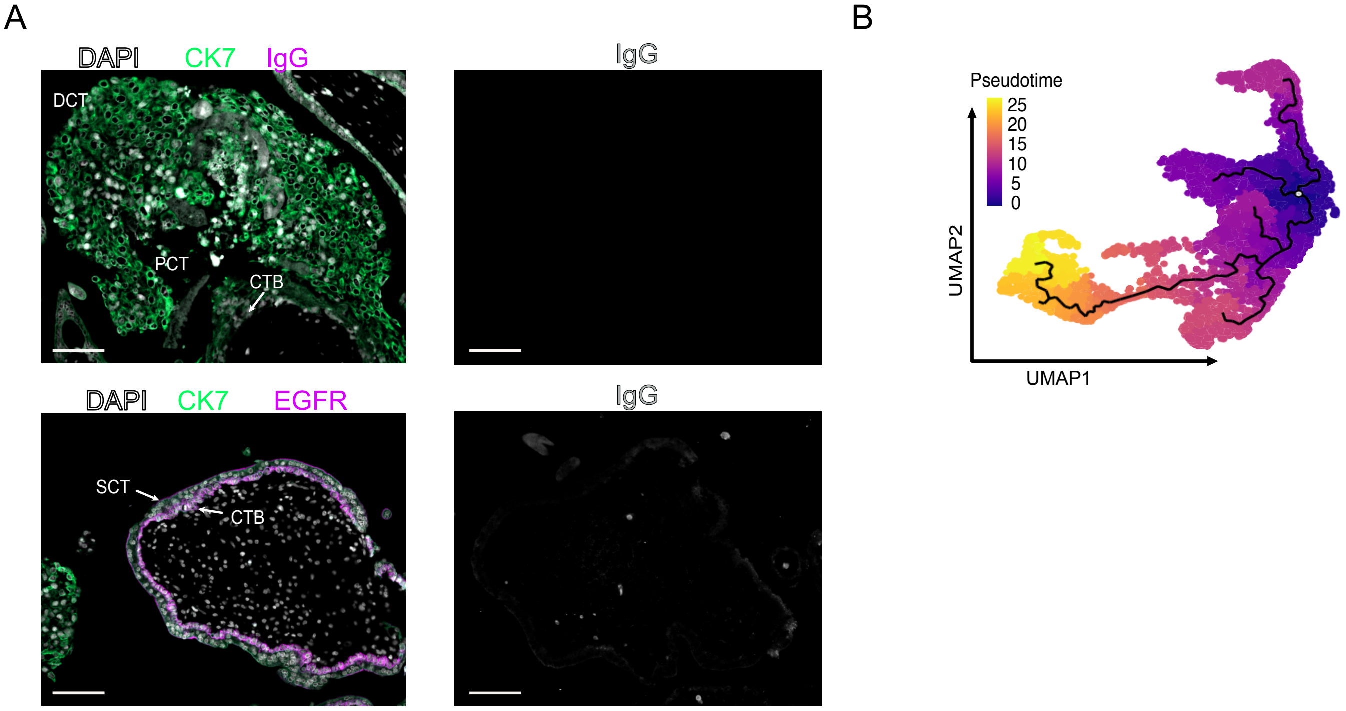

### Supplemental Figure 3

#### Supplemental Figure 3

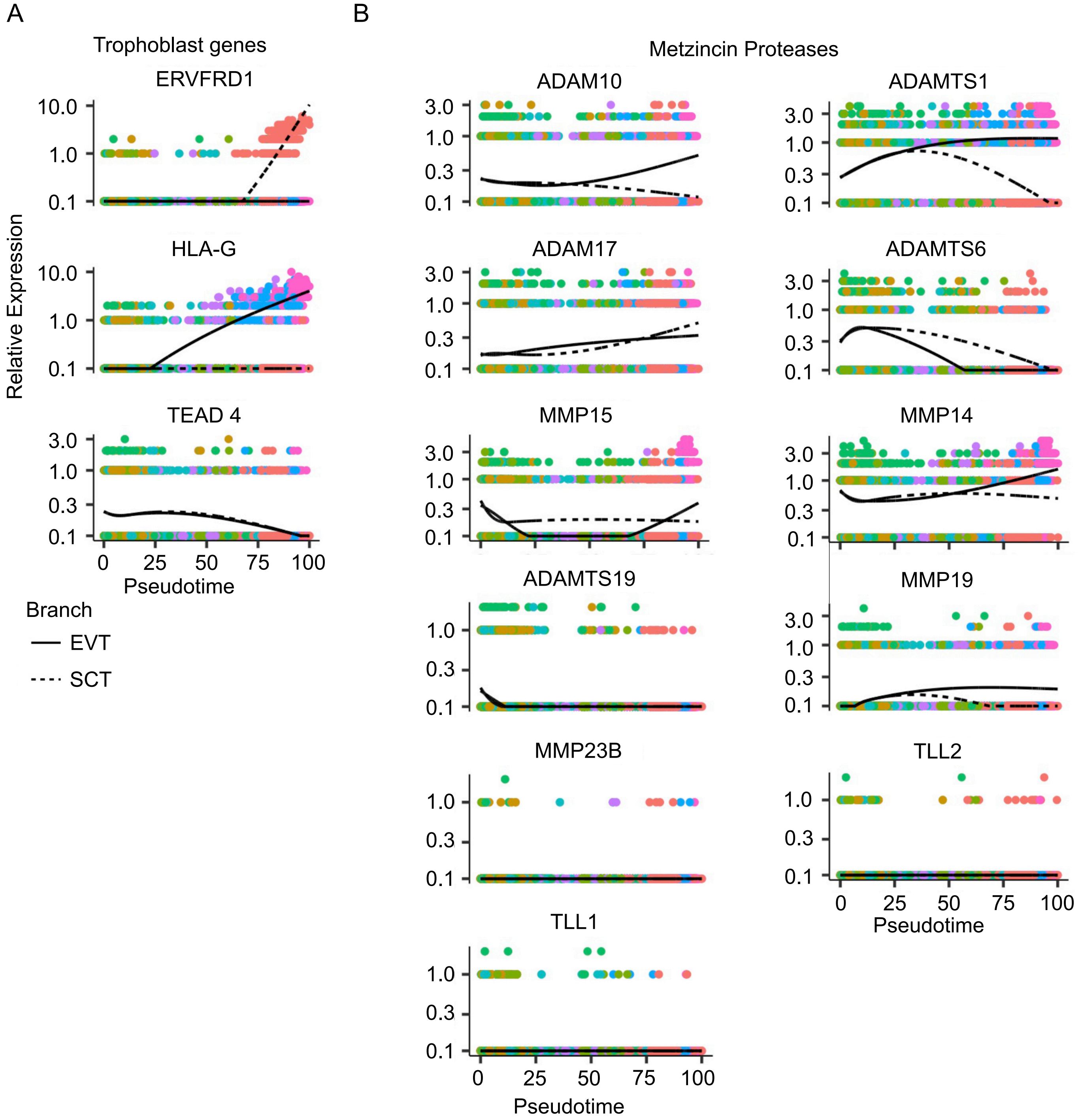

### Supplemental Figure 4

##### Supplemental Figure 4

A

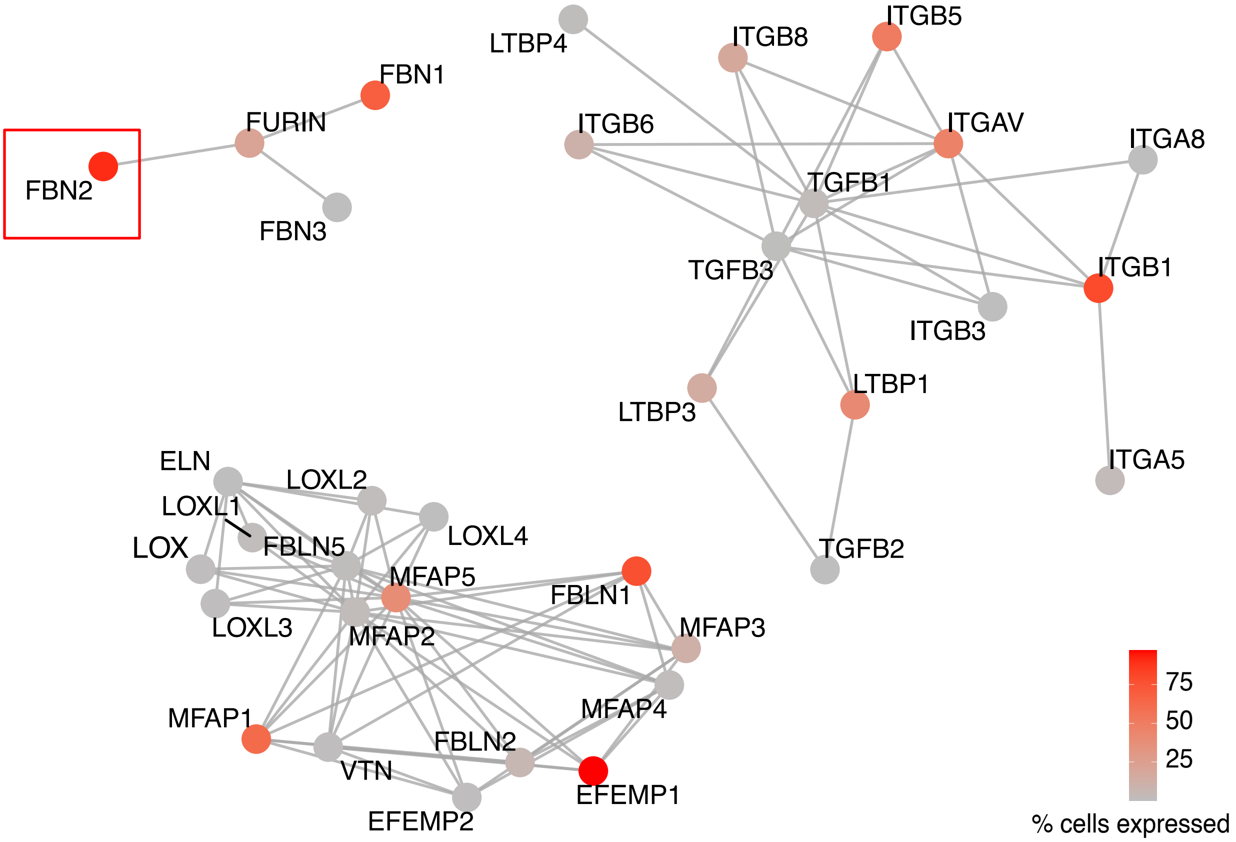
